# Water Resource Management: IWRM strategies for Improved Water Management. A Systematic Review of Case Studies of East, West and Southern Africa

**DOI:** 10.1101/2020.07.17.208413

**Authors:** Tinashe Lindel Dirwai, Edwin Kimuta Kanda, Aidan Senzanje, Toyin Isiaka Busari

## Abstract

**Objective:** The analytical study systematically reviewed the evidence about the IWRM water strategy model. The study analysed the IWRM strategy advances and practical implications it had, since inception on effective water management in East, West and Southern Africa.

**Methods:** The study adopted the Preferred Reporting Items for Systematic Review and Meta-analysis Protocols (PRISMA-P) and the scoping literature review approach. The study searched selected databases for peer-reviewed articles, books, and grey literature. DistillerSR software was used for article screening. A constructionist thematic analysis was employed to extract recurring themes amongst the regions.

**Results:** The systematic literature review detailed the adoption, policy revisions and growing/emerging policy trends and issues (or considerations) on IWRM in East, West and Southern Africa. Thematic analysis derived four cross-cutting themes that contributed to IWRM strategy implementation and adoption. The identified four themes were donor effect, water scarcity, transboundary water resources, and policy approach. The output further posited questions on the prospects, including whether IWRM has been a success or failure with the African water resource management fraternity.

**Graphical Abstract:** 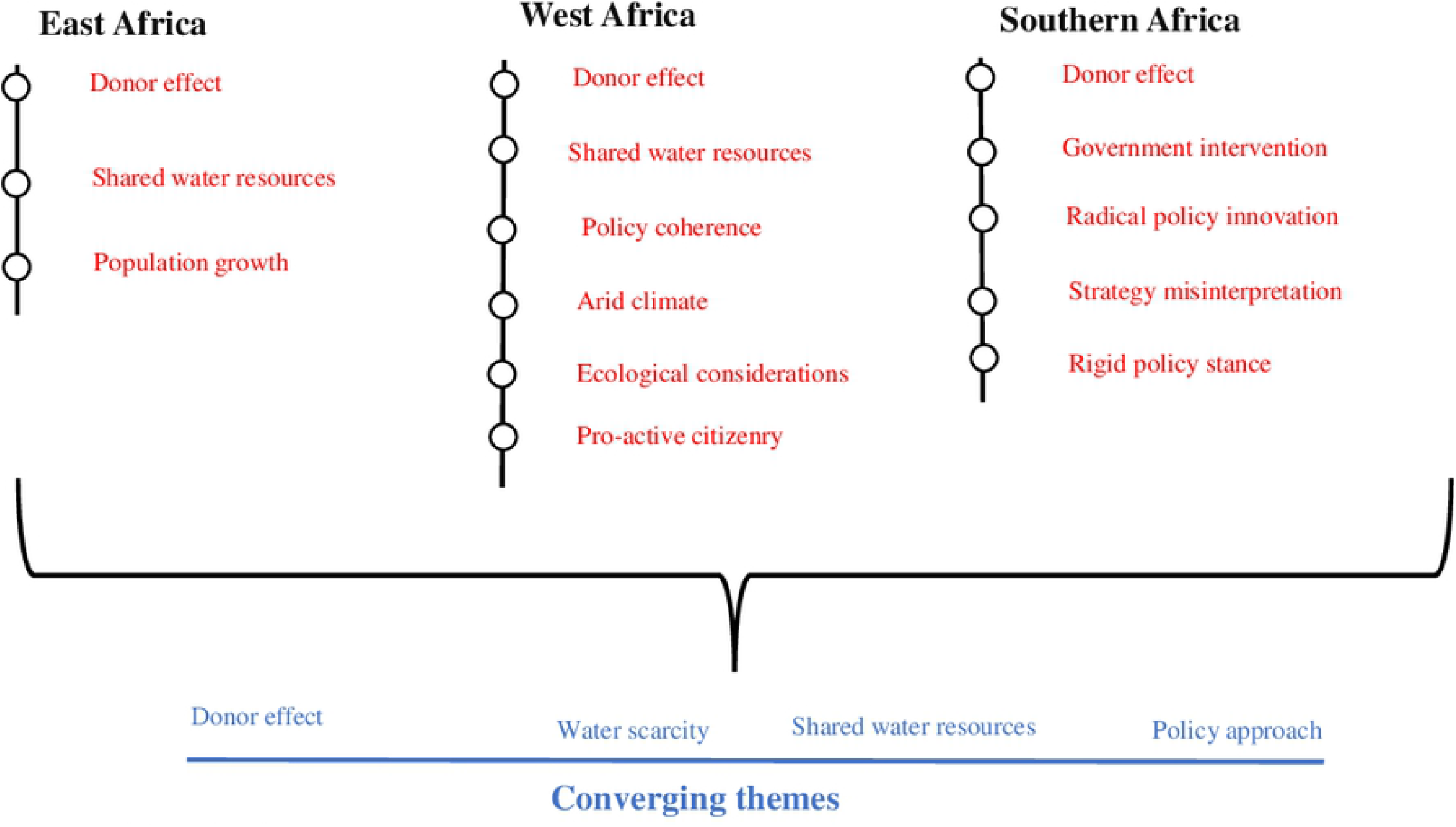

## 1 Introduction

Integrated Water Resources Management (IWRM) is a concept that is meant to foster effective water resource management. GWP [1] defined it as “the process which promotes the coordinated development and management of water, land and related resources, to maximise the resultant economic and social welfare equitably without compromising the sustainability of vital systems”. A holistic approach, in the form of the Dublin statement on Water and Sustainable Development (DSWSD), emerged and it became the backbone of IWRM principles.

IWRM strategy approach and implementation are ideally linked to individual countries developmental policies [2]. Southern Africa (Zimbabwe and South Africa) is the biggest adopter of the water resource management strategy and produced differed uptake patterns [3]. Tanzania benefited from donor funds and World Bank programmes that sought to alleviate poverty and promote environmental flows. The World Bank radically upscaled and remodelled IWRM in Tanzania through the River Basin Management – Smallholder Irrigation Improvement Programme (RBM-SIIP) [4].

Uganda latently adopted the IWRM strategy in 1992 [5]. Government of Uganda’s efforts of liberalising the markets, opening democratic space and decentralising the country attracted donor funds that drove the IWRM strategy agenda. The resultant effort led to the formation of the Nile Basin Initiative (NBI) [5, 6]. The long-standing engagement between Uganda and the Nordic Fresh Water initiative helped in the diffusion of IWRM in the country. Burkina Faso and Ghana made significant strides in operationalising the IWRM policy approach by adopting the West Africa Water Resources Policy (WAWRP). A massive sense of agency coupled with deliberate government efforts drove the adoption status of Burkina Faso.

Total innovation diffusion can be achieved when the practice or idea has supporting enablers. Innovation policy is key in altering societal orthodox policy paths that fuel hindrance and subsequently in-effective water governance [7]. Acknowledging the political nature of water (water governance and transboundary catchments issues) is the motivation to legislate water-driven and people-driven innovative policy [8]. Water policy reform should acknowledge the differing interests’ groups of the water users and its multi-utility nature; thus, innovation diffusion channels should be tailored accordingly, avoiding the ‘one size fits all’ fallacy.

IWRM as an innovative strategy approach diffused from the global stage to Africa and each regional block adopted the approach at different times under different circumstances.

The paper sought to unveil the innovative IWRM strategy approach by critically examining its genesis, implementation, adoption and the main drivers in smallholder farming fraternity. The study was informed by Flows and Practices. The Politics of IWRM in Africa.

## 2 Conceptual Framework and Methodology

The analytical framework applied in the study (Figure 1) is based on the water innovation frames by the United Nations Department of Economic and Social Affairs [9]. The UNDESA Γ9ļ, classified water frames into three distinct categories namely water management strategies e.g., IWRM, water infrastructure and water services. The former partly involves IWRM strategies and the latter encompasses economic water usage such as agriculture, energy production and industrial applications [8]. The review also adopted the thematic analysis approach by Braun and Clarke [10] to extract, code, and select candidate converging themes for the systematic review.

**Figure 1.**
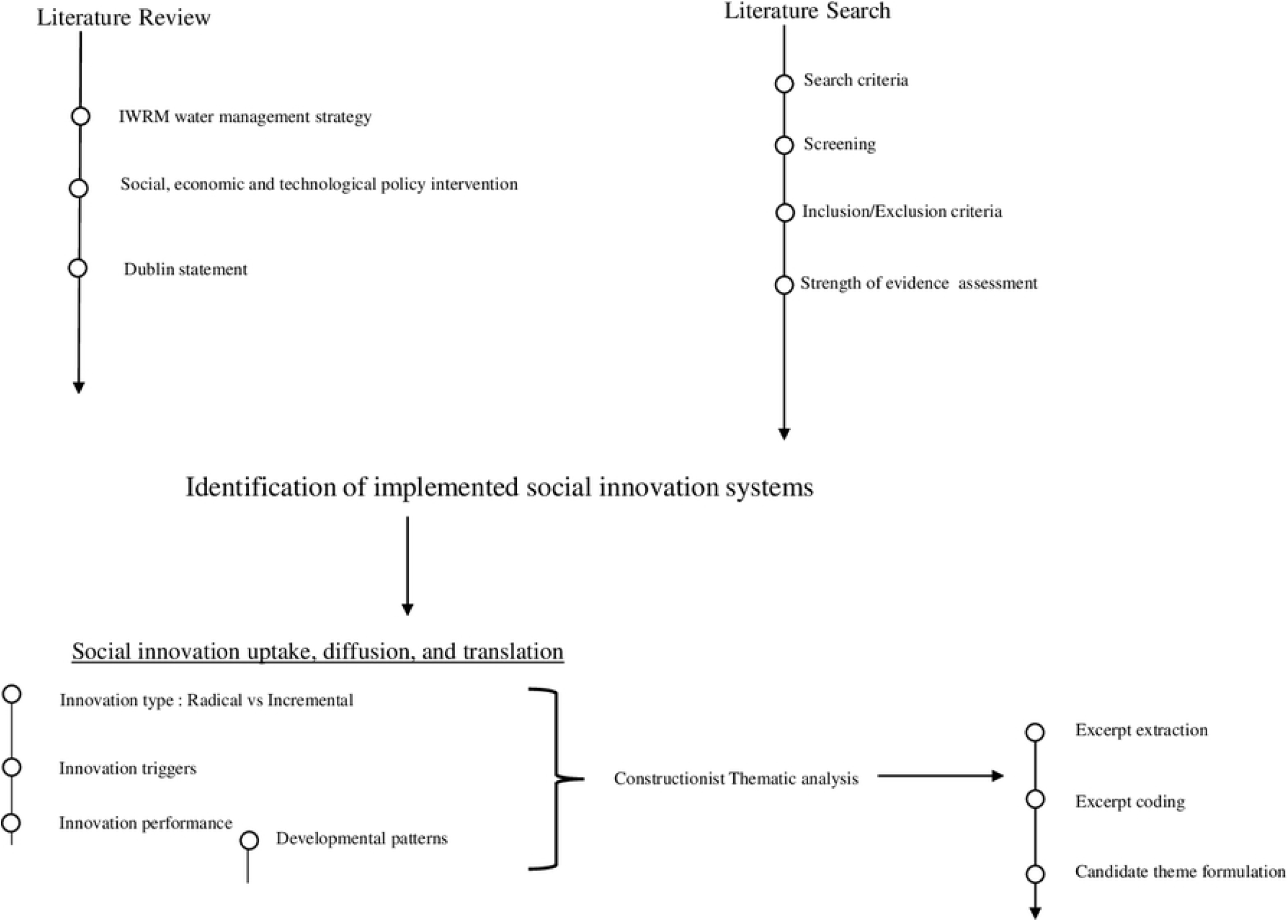
The water framing, social innovation diffusion, and constructionist thematic conceptual framework (source: Authors)

The literature review identified research gaps that informed the employed search strategy. The literature that qualified for inclusion was thoroughly analysed and discussed. The aggregated outcomes were used excerpt extraction in the thematic analysis.

### 2.1 Methodology

The study adopted the Preferred Reporting Items for Systematic Review and Meta-Analysis Protocols (PRISMA-P) and the <u>Arksey and O’Malley</u> [11] methods to scope, gather, screen and report literature. This section follows the systematic literature review framework by <u>Moher, Shamseer</u> [12] which yields accurate and unbiased evidence-based conclusions [13].

#### 2.1.1 Eligibility criteria

For information gathering the study searched the following databases: Scopus, Web of Science, Google Scholar, UKZN-EFWE, CABI, JSTOR, African Journals Online (AJOL), Directory of Open Access Journals (DOAJ), J-Gate, SciELO, WorldCat, WorldWideScience and AgeLine for peer-reviewed articles, books, and grey literature. The study did not emphasize publication date as recommended by <u>Moffa, Cronk</u> [14].

#### 2.1.2 Search strategy

The search strategy or query execution [13] utilised Boolean operators (**OR** & **AND**). The dynamic nature of the search strategy required the authors to change the search terms and strategy, for example, if digital databases did not yield the expected search items the study would manually search for information sources. The search queries included a string of search terms i.e., IWRM, water management - *East Africa* - *West Africa* **AND/OR** - *Southern Africa*.

#### 2.1.3 Selection process

DistillerSR© software was used for article screening. The screening was based on the article title, abstract and locality. The study employed a two-phase screening process [15], the first phase screened according to title and the second phase screened according to abstract and keywords. An inclusion-exclusion criterion was established for streamlining literature and quality check.

#### 2.1.4 Strength of Evidence

The study employed a strength of evidence exercise [16]. This was done to validate the literature sources. The exercise adopted a four-level assessment approach to validate the relevance of the included literature for analysis (Table 1). The grading follows that substantiated and documented research is given a *high* score, whereas a *moderate* score is assigned to partially substantiated studies and studies with a conditional conclusion. Opinion papers are assigned a *low* score and none-evidenced based researches are ungraded.

**Table 1.**
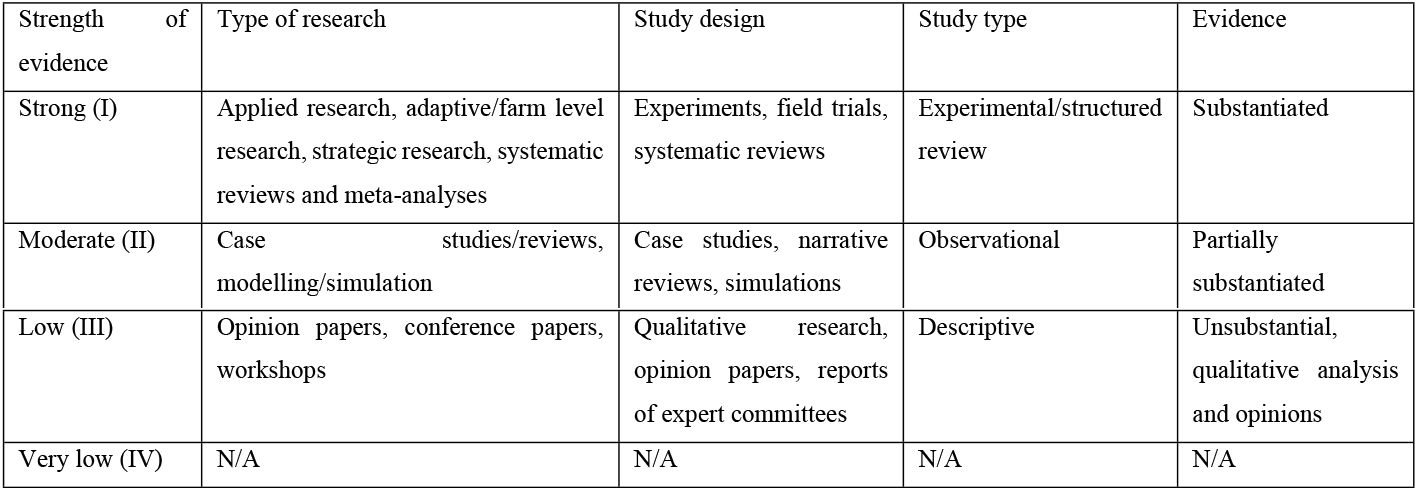
Strength of evidence grading.

## 3 Results and Discussion

Data charting comprised of the PRISMA flow-chart (Figure 2). The study utilised 70 out of 183 records (n= 37, 46%) for East Africa, (n= 37, 46%) for Southern Africa, and (n= 6, 8%) for West Africa.

**Figure 2.**
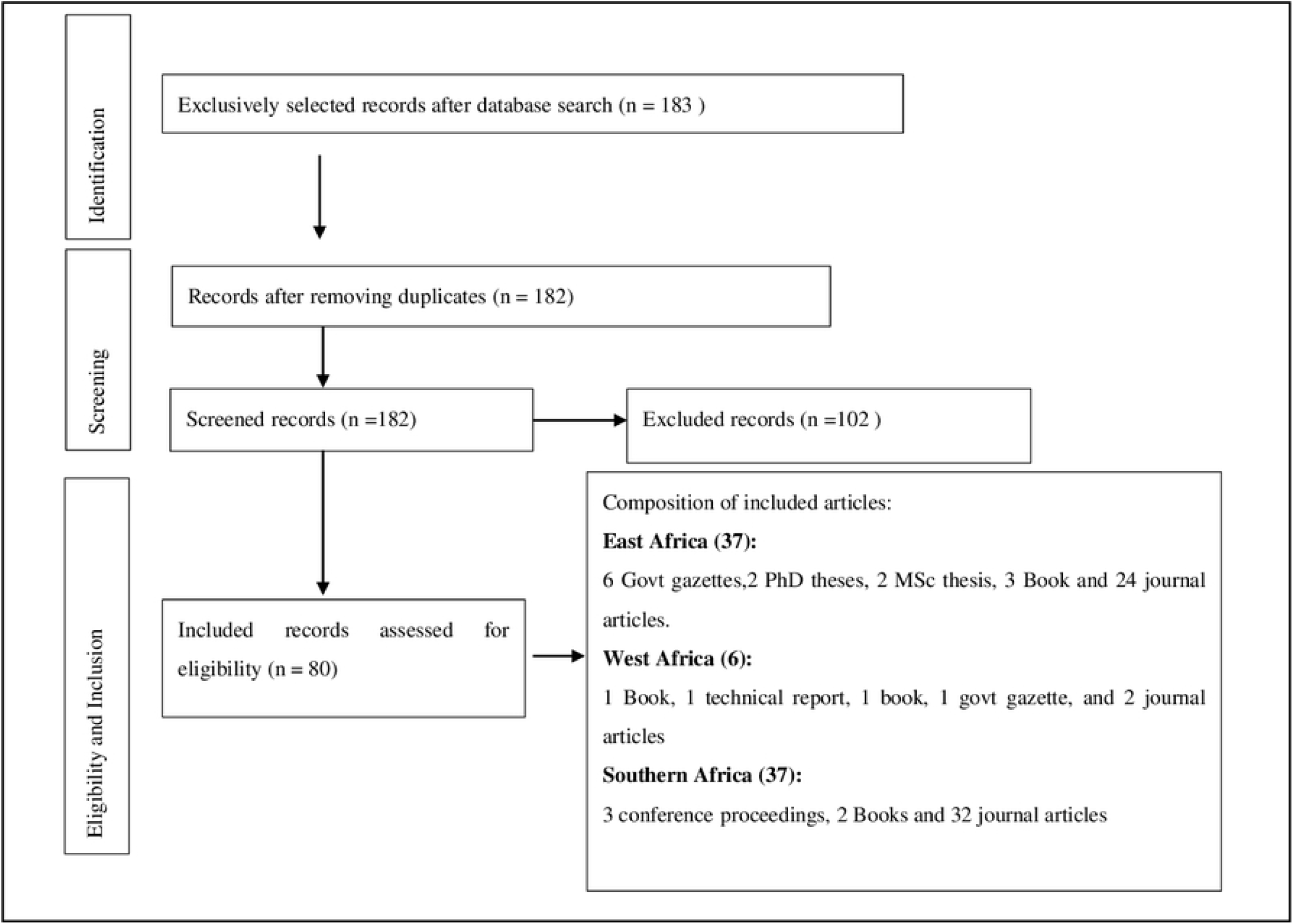
Systematic review flowchart based on PRISMA flowchart [12]

The relevant studies were charted and graded according to <u>Thomas, Georgios</u> [16] grading method (Table 2).

**Table 2.**
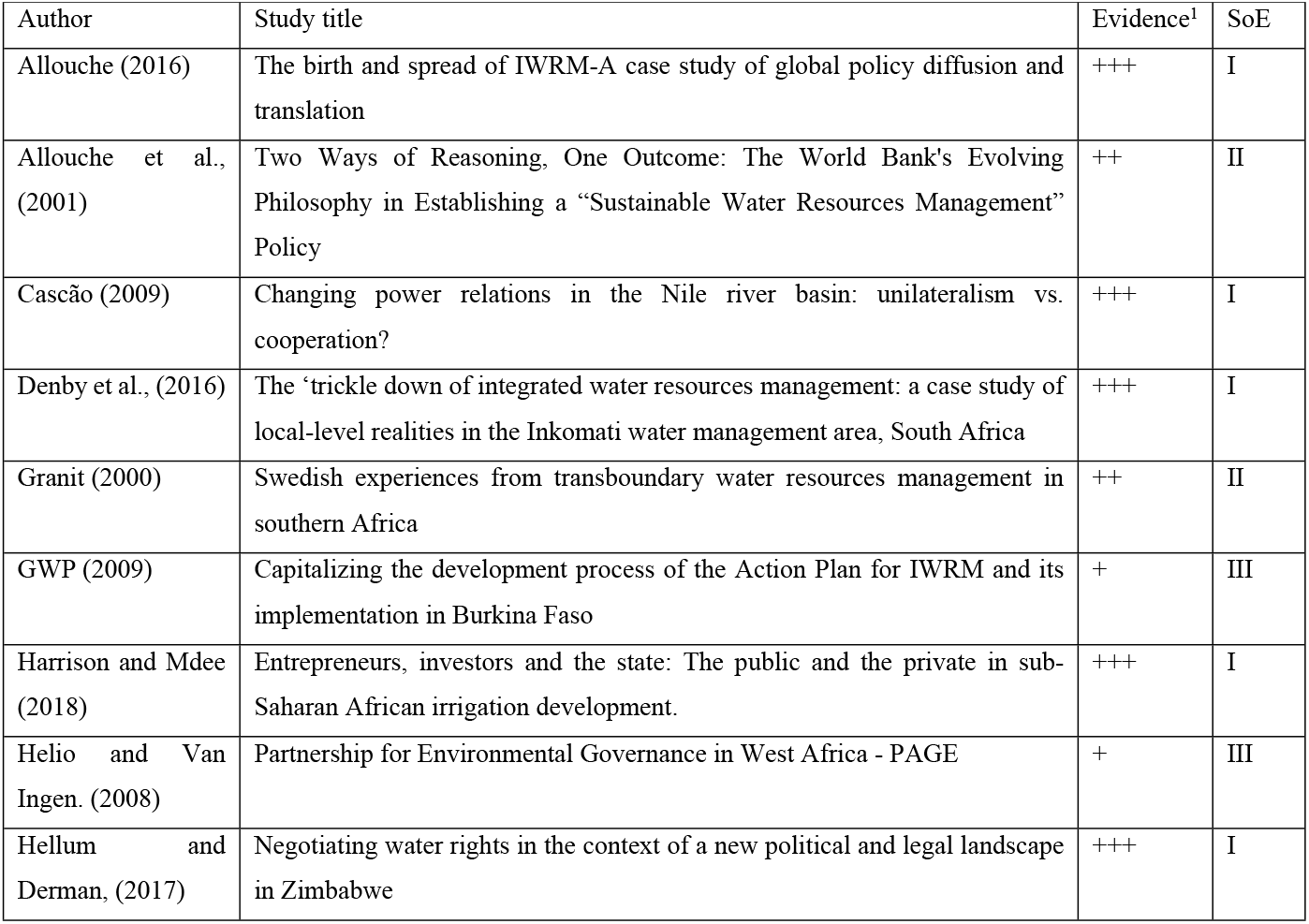
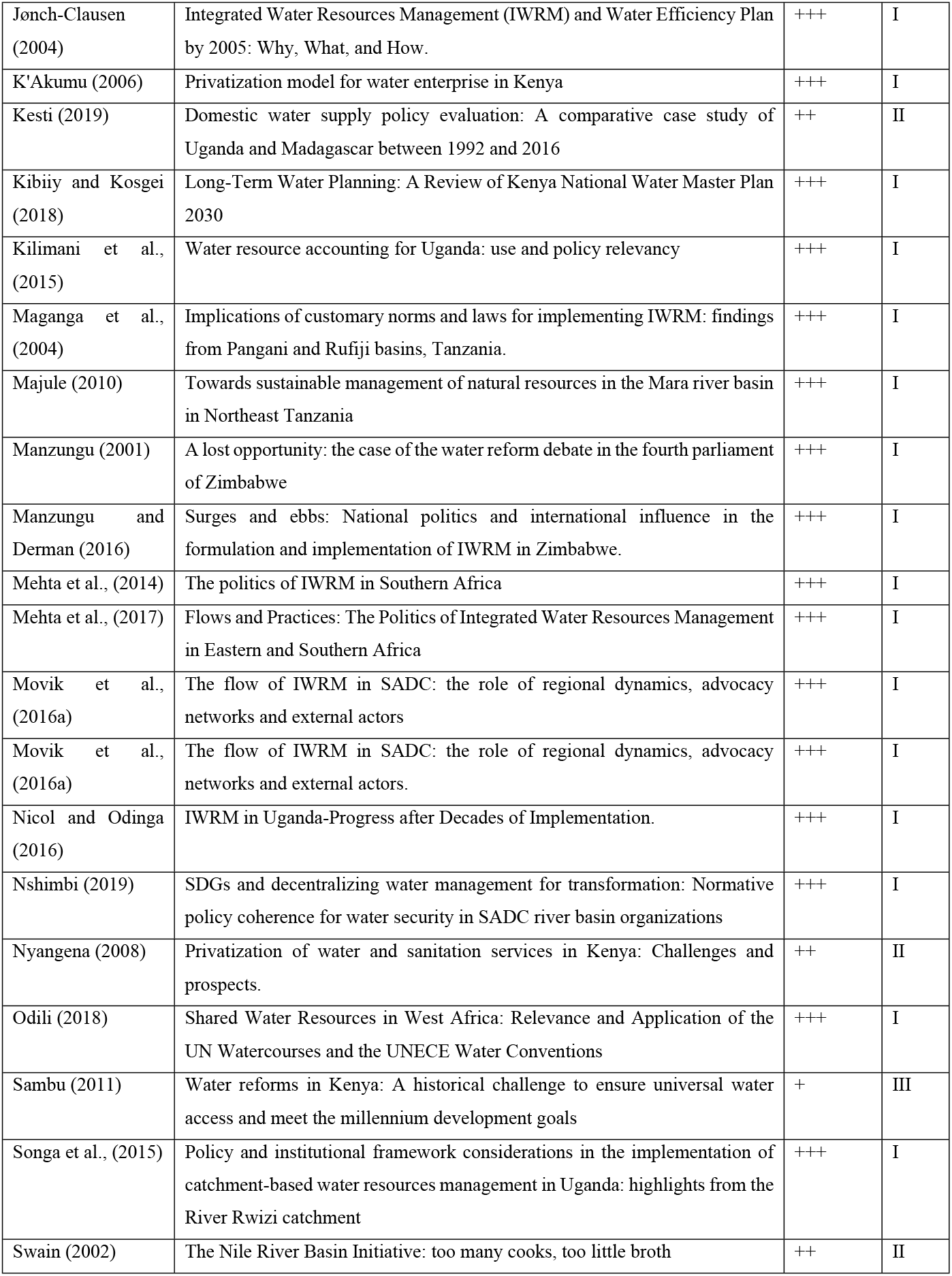
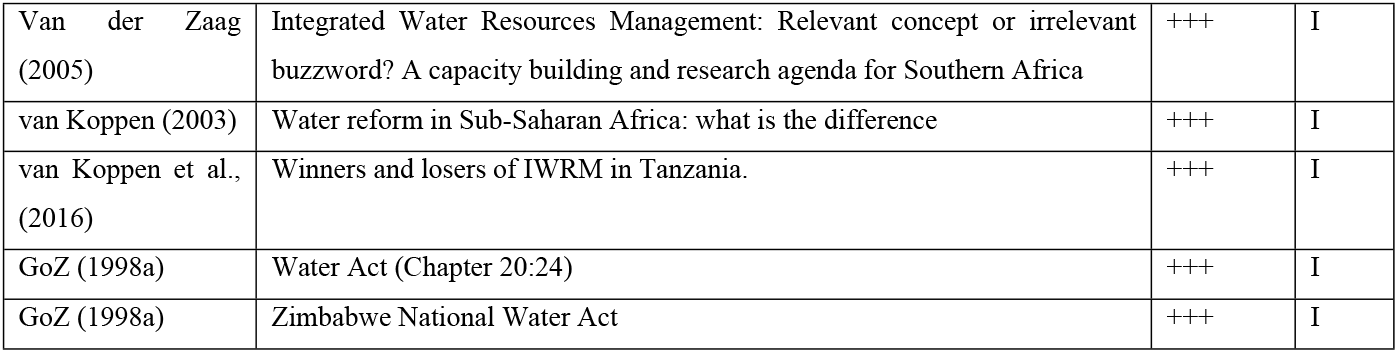
Grading results for the selected literature.

## 4 Case Studies

### 4.1 East Africa

The introduction of IWRM in the region was initiated in 1998 by the water ministers in the Nile basin states due to the need for addressing the concerns raised by the riparian states. These water sector reforms revolved around the Dublin principles initiated by the UN in 1992 [17]. In 1999, Kenya developed the national water policy and the enabling legislation, the Water Act 2002 was enacted [18]. The Act was replaced by the Water Act 2016 which established the Water Resources Authority (WRA) as the body mandated to manage water resources in line with the IWRM principles and Water Resource Users Association (WRUA) as the lowest (local) level of water management [19].

Similarly, Uganda developed the national water policy in 1999 to manage, and develop the available water resources in an integrated and sustainable manner [20]. The National Water Policy further provides for the promotion of water supply for modernized agriculture [21]. Tanzania’s water policy of 2002 espouses IWRM principles, and its implementation is based on a raft of legal, economic, administrative, technical, regulatory and participatory instruments [22]. The National Irrigation Policy (NIP), 2010 and the National Irrigation Act, 2013 provides the legal basis for the involvement of different actors on a private-public partnership basis [23].

#### 4.1.1 Diffusion innovation drivers of IWRM in East Africa

##### (i) Water Scarcity

The adoption of IWRM was necessitated by water scarcity which is experienced by the countries in the region, which formed the need for adoption of prudent water resources management strategies as envisaged under the Dublin principles which was championed indirectly, according to <u>Allouche</u> [24], by the World Bank. Specifically, the need to give incentives and disincentives in water use sectors to encourage water conservation.

Kenya is a water-scarce country with per capita water availability of 586 m^3^ in 2010 and projected to 393 m^3^ in 2030 [25]. Uganda is endowed with water resources, however, it is projected that the country will be water-stressed by 2020 which could be compounded by climate variability and change, rapid urbanization, economic and population growth [26].

Using water scarcity was in essence coercing countries to adopt the IWRM principles with the irrigation sector, the contributor of the largest proportion of water withdrawals, becoming the major culprit [24]. The researchers opine that the effects of water scarcity in the region can be countered by adopting IWRM policy approach, butadaptively to suit the local context and thus, persuasive rather than coercive, is the appropriate term. Indeed, as put forward by Van der Zaag [27], IWRM is not an option but it is a must and therefore, countries need to align their water policies and practices in line with it.

##### (ii) Trans-boundary Water Resources

Water resources flow downstream indiscriminately across villages, locations, regions and nations/states and therefore necessitates co-operation. The upstream and downstream relationships among communities, people and countries created by the water is asymmetrical in that the actions upstream tend to affect the downstream riparian and not the other way round [27]. In East Africa, the Nile Basin Initiative (NBI) and the Lake Victoria Basin Commission (LVBC) plays a critical component in promoting the IWRM at regional level [17].

The Nile River system is the single largest factor driving the IWRM in the region. Lake Victoria, the source of the Nile River is shared by the three East African states of Kenya, Uganda and Tanzania. Irrigation schemes in Sudan and Egypt rely exclusively on the waters of River Nile and are therefore apprehensive of the actions of upstream states notably Ethiopia, Kenya, Uganda, Tanzania, Rwanda and Burundi. The source of contention is the asymmetrical water needs and allocation which was enshrined in the Sudan-Egypt treaty of 1959 [28]. All the riparian countries in the Nile basin have agricultural-based economies and thus irrigation is the cornerstone of food security [29]. Therefore, there was the need for the establishment of basin-wide co-operation which led to the formation of NBI in 1999 with a vision to achieve sustainable socio-economic development through the equitable utilisation of the Nile basin water resources [30].

The Mara River is another trans-boundary river which is shared between Tanzania and Kenya and the basin forms the habitat for the Maasai Mara National Reserve and Serengeti National Park in Kenya and Tanzania, respectively, which is prominent for the annual wildlife migration. Kenya has 65% of the upper part of the basin, any development on the upstream, such as hydropower or water diversion, will reduce the water quantities and therefore affect the Serengeti ecosystem and the livelihoods of people in Tanzania [31]. The LVBC, under the East African Community, developed the Mara River Basin-wide – Water Allocation Plan (MRB-WAP) to help in water demand management and protection of the Mara ecosystem [32]. The mandate of the LVBC is to implement IWRM in Lake Victoria Basin riparian countries [17].

Other shared water basins include the Malakisi-Malaba-Sio River basin shared between Uganda and Kenya and the Kagera River basin traversing Burundi, Rwanda, Tanzania and Uganda. The two river basins form part of the Upper Nile system and are governed through the LVBC and the NBI.

##### (iii) Donor Influence

The World Bank has been pushing for IWRM principles through the NBI and by pressurising Egypt to agree to co-operate with the upstream riparian countries in the Nile basin [29]. In the early 1990s, the World Bank had aligned its funding policies to include sustainable water resources management [33].

Privatization was part of the Structural Adjustment Programmes (SAPs) introduced by the World Bank and International Monetary Fund (IMF) as part of ensuring the sustainability of public enterprises which were bogged down by inefficiencies of state-centred approach to management and these two institutions advocated for government withdrawal in the affairs of these entities [34]. In Kenya, the World Bank was the one identifying the firms to be privatized which were mostly in the electricity, telecommunication and the water sectors [35].

In Tanzania, Norway, through NORAD, played a key role in implementing IWRM by promoting water projects including hydropower schemes [36]. Indeed the transformation of the agricultural sector in Tanzania through *Kilimo Kwanza* policy of 2009 which emphasises on the commercialization of agriculture including irrigation was driven by foreign donors such as the USAID and UK’s DFID [23].

In Uganda, however, the reforms in the water sector were initiated devoid of external influence [37]. However, this assertion is countered by Allouche [24] who pointed that Uganda had become a ‘darling’ of the donor countries in the early 1990s and that DANIDA helped to develop the Master Water Plan and the country was keen to show a willingness to develop policy instruments favourable to the donor. East African countries are developing economies and therefore most of their development plans are supported by external agencies, which to some extent come with subtle ‘conditions’ such as free-market economies. In fact imposition of tariffs and other economic instruments used to implement IWRM in water supply and irrigation is a market-based approach which was favoured by the World Bank and other development agencies.

The three East Africa countries are at various stages of implementation of IWRM, as illustrated in Table 3, and have made substantial progress and achievements in enacting policies and legislations and institutional set-up regarding IWRM. Besides, the countries, except for Tanzania, have made substantial achievements in building institutional capacity. However, all the countries have made little progress in infrastructure and sustainable financing [17].

**Table 3.** Status of IWRM implementation [38].

| Country | Level | Interpretation of status |
| --- | --- | --- |
| Kenya | Medium - high | Capacity to implement IWRM elements is generally adequate and elements are generally being implemented under long-term programmes |
| Uganda | Medium-high | Capacity to implement IWRM elements is generally adequate and elements are generally being implemented under the long-term programme |
| Tanzania | Medium-low | IWRM elements are generally institutionalized and implementation is underway. |

Kenya’s water and sanitation are governed by the Water Act, 2016 while irrigation is under the Irrigation Act, 2019. The Irrigation Act established the National Irrigation Authority (NIA) which is mandated to develop irrigation infrastructure for national and public schemes and provide extension services for private and smallholder schemes, [39]. Separated irrigation and water and sanitation sector, in the researchers’ opinion, leads to a fragmented approach to water resources management which hinder innovation diffusion. <u>Sambu</u> [40], argued under this arrangement, concerns of water quality, quantity and sanitation are handled differently leading to disjointed and uncoordinated interventions.

Similarly, in Uganda, irrigation is domiciled in the Ministry of Agriculture, Animal Industry & Fisheries (MAAIF) while the Ministry of Water and Environment (MWE) deals with water supply, water management and sanitation where the IWRM concept is implemented, which may lead to duplication and overlaps [41]. Although MWE has an inter-ministerial framework which allows it to link with other ministries, such as MAAIF with respect to irrigation, there is lack of clarity which leads to overlaps and conflicts with other ministries [21].

Failure to acknowledge customary systems hinders IWRM implementation in smallholder irrigation in East Africa. For instance, in the Rufiji and Pangani river basins in Tanzania, the water rights system is not well defined, is poorly implemented and fails to incorporate customary arrangements on water allocation [42]. An incongruous water permit and customary system impact negatively on the operationalisation of the IWRM policy approach [43].

#### 4.1.2 Prospects of IWRM in East Africa

The implementation of IWRM in the region, and more so the irrigation sub-sector, will continue to evolve amid implementation challenges. The dynamics of water policies, increased competition for finite water resources from rapid urbanization, industrialization and population growth will continue to shape IWRM practices in the region. Trans-boundary water resources management will possibly take centre stage as East African countries move towards full integration and political federation as envisaged in the four pillars of the East African Community treaty.

Kenyan water bodies transcend across Counties and the attempt to align the water law with the Constitution which culminated in the Water Act, 2016 is still facing implementation challenges. The County Governments and the National government through national institutions such as the NIA, WRA and the Ministry of water and irrigation need to develop a working framework for water resource management, water supply and sanitation. Decision support tools will be very relevant in the trans-boundary water resources such as the Nile system, Mara and Kagera river basins.

### 4.2 West Africa

West Africa possesses an unregistered IWRM policy approach that is espoused in the West Africa Water Resources Policy (WAWRP) of 2008. The WAWRP is founded on the following legal principles; (a) “promote, coordinate and ensure the implementation of a regional water resource policy in West Africa, in accordance with the mission and policies of ECOWAS” and (b) “harmonization and coordination of national policies and the promotion of programmes, projects and activities, especially in the field of agriculture and natural resources”. The founding legal basis resonates with the Dublin principles.

The WAWRP design actors were the Economic Community of West African States (ECOWAS), Union Economique et Monétaire Ouest Africaine (UEMOA), and Comité Permanent Inter-État de Llutte Contre la Sécheresse au Sahel (CILSS). CILSS CILSS is the technical arm of ECOWAS and UEMOA. The institutional collaboration was driven by the fact that West Africa needed a sound water policy for improved regional integration and maximised economic gains. ECOWAS established the Water Resources Coordination Centre (WRCC) to (a) oversee and monitor the region’s water resources and management activities and (b) to act as an executive organ of the Permanent Framework for Coordination and Monitoring (PFCM) of IRWM [44].

The inception and triggers of IWRM in West Africa can be traced back to the General Act of Berlin in 1885 which, among other things, dictated water resources use of the Congo and Niger rivers [45]. A multiplicity of agreements around shared watercourses in West Africa led to the realisation of the IWRM policy approach. For example, the Senegal River Basin (SRB) Development Mission facilitated collaboration between Senegal and Mauritania in managing the SRB. Another noteworthy agreement was Ruling C/REG.9/7/97, a regional plan to fight floating plants in the ECOWAS countries [45].

#### 4.2.1 Diffusion innovation drivers of IWRM in West Africa

GWP (2003) categorised the countries according to the level of adoption into three distinct groups namely; (a) Group A comprises of countries with the capacity to develop and adopt the IWRM approach (Burkina Faso and Ghana), (b) Group B comprises of countries needing “light support” to unroll the IWRM plan (Benin, Mali, Nigeria, and Togo), and (3) Group C comprises of laggards, needing significant support to establish an IWRM plan (Cape Verde, Ivory Coast, Gambia, Guinea, Guinea Bissau, Liberia, Mauritania, Niger, Senegal and Sierra Leone).

##### (i) Water Scarcity

West African climatic conditions pose a threat on the utilisation of the limited water resource. Utilisation is marred by erratic rainfalls and primarily a lack of water resources management know-how [44]. Countries in the Sahelian regions are characterised by semi-arid climatic conditions. Thus, dry climatic conditions account as an innovation driver to ensure maximised water use efficiency. Although the region acknowledges the need for adopting the IWRM policy approach, they have varied adoption statuses (GWP, 2003).

##### (ii) Government Intervention and Regional Bloc Pro-Activity

The Burkinabe government exhibited political goodwill such that in 1995 the government brought together two separate ministries into one ministry of Environment and Water thus enabling coherent policy formulation and giving the ministry one voice to speak on water matters.

The dynamic innovation arena (where policy players interact) allows continuous policy revision and redesign thus water policy reform innovation diffusion, and policy frameworks are in a perpetual state of shifting. For example, in the 1990s the Burkinabe government was engaged in several water-related projects and was continuously experimenting with local governance and privatization (from donors) [1]. This policy shift according to <u>Gupta</u> [46] qualifies as an innovation driver.

##### (iii) Trans-boundary Water Resources

The universal transboundary nature of water creates dynamics that warrant cooperation for improved water use. West Africa has 25 transboundary watercourses and only 6 are under agreed management and regulation. The situation is compounded by the fact that 20 watercourses lack strategic river-basin management instruments [45]. Unregistered rules and the asymmetrical variations associated with watercourses warrant the introduction of the IWRM principle to set equitable water sharing protocols and promote environmental flows (eflows). The various acts signed represent an evolutionary^2^ treaty development that combines the efforts of riparian states to better manage the shared water resources.

##### (iii) Donor Influence

Donor aid cannot be downplayed in pushing for policy diffusion in low-income aid-dependent countries. <u>GoBF</u> [47] cites that from the period 1996 – 2001, more than 80% of water-related projects were donor funder. Research by <u>Cherlet and Venot</u> [48] also cites that almost 90% of the water investments in Mali were funded outside the government apparatus. It can be argued that donor-aid plays a pivotal and central role in diffusing policy and innovation in aid-depended countries.

##### (iv) Pro-active Citizenry

Burkina Faso and Mali’s adoption story is accentuated by heightened agency, the individual enthusiasm on influencing the outcome facilitated policy diffusion and can be argued to be a potential innovation diffusion driver for the IWRM policy approach in the region. The individual policy diffusion fuelled by an enthusiastic citizenry was a sure method that effectively diffused awareness around the IWRM innovation and acted as a driver of the IWRM practices in the region. Individual strategies were honed in smallholder farming institutions to diffuse the IWRM practice and drawing from the <u>Sabatier and Jenkins-Smith</u> [49] advocacy coalition theory, having individuals with common agendas promoted the transfer and diffusion of water reforms in parts of West Africa.

Adoption of the IWRM policy in West Africa is fraught with many challenges. For example, despite having significant water resources, the lack of a collective effort by the governments to train water experts at national level presents a challenge for adoption. Unavailability of trained water experts (who in any case are diffusion medium) results in a lack of diffusion channels that facilitate policy interpretation, translation and its subsequent implementation. <u>Helio and Van Ingen</u> [44] pointed out how political instability possesses a threat to current and future implantation initiatives.

#### 4.2.2 Prospects of IWRM in West Africa

The future collaboration projects and objective outlined by ECOWAS, CILSS, and UEMO highlight a major effort to bring the region to speed with the IWRM policy approach. The WAWRP objectives can potentially set up the region on an effective IWRM trajectory which can be mimicked and upscaled in other regions. Positives drawn from the region are the deliberate institutional collaborations. Burkina Faso and Mali have the potential to operationalise and facilitate policy diffusion to other neighbouring states. Donor driven reform is essential and national ownership is critical in ensuring the water reform policies and innovation diffusion processes are implemented at the national level.

### 4.3 Southern Africa

Southern Africa has over 15 shared transboundary river basins. SADC member states established the Protocol on Shared Water Systems (PSWS) which meant to encourage sustainable water resources utilisation and management. The PSWS was perceived to strengthen regional integration [50]. The regional bloc formulated the Regional Strategic Action Plans (RSAPs) that sought to promote an integrated water resources development plan. The action initiative mimicked IWRM principles and the cascading shared water resources initiatives acted as a catalyst for the genesis of IWRM in Southern Africa [43]. SADC houses the Waternet and the GWP-SA research and innovation hubs upon which SADC’s adoption was anchored on. Besides, the availability of trained water experts in the region who were willing to experiment with the IWRM policy approach and water scarcity fuelled by climate change prompted the region’s adoption of the IWRM policy approach at the local level

#### 4.3.1 Diffusion innovation drivers and adoption status in Southern Africa

This section will analyse two case studies from the region, i.e. Zimbabwe and South Africa. The case studies analyse the triggering factors in each country and the practical implication on the local level realities.

##### 4.3.1.1 Zimbabwe

In 1998 a National Water Policy (NWP) which was in alignment with the Dublin principles was enacted, thus giving Zimbabwe an IWRM policy footing. The NWP explicitly acknowledged that; water was a finite resource and it needed conservation, and water was a commodity for economic good, a clear commitment to the IWRM [3].

###### (i) Fast Track Land Reform (FTLR) Programme and Institutional Incoherence

The Fast Track Land Reform (FTLR) programme disaggregated the large-scale commercial farms and created smallholder farming [51], consequently influencing and dictating IWRM policy path. The FTLR programme had a negative impact on the spread and uptake of IWRM. A series of poor economic performance and poor policy design compounded the limited diffusion and the adoption of IWRM practices at local levels in Zimbabwe. The FTLR programme compounded the innovation diffusion process as the Zimbabwe National Water Authority (ZINWA) lost account of who harvested how much at the newly created smallholder farms. Thus, water access imbalance ensured, and ecological sustainability was compromised.

Policy incoherence was a major factor in poor IWRM diffusion and adoption, for example, the government did not synchronise the land and water reforms thus it meant at any given point in time there was a budget for one reform agenda [3] and the land reform agenda would take precedence because of political rent-seeking.

###### (ii) Donor Aid

Whilst the World Bank and western donor organisations funded the water-related projects in the urban setting there was no budget earmarked for smallholder farming and IWRM-related activities^3^ [3]. A lack of access to international funding and fleeting donor aid exacerbated the policy uptake as such the anticipated implementation, operationalisation and continuous feedback mechanism for policy revision and administering process was never realised. Continued clashes of water and land reforms have created laggards in the IWRM adoption.

###### (iii) Lack of Politics of Pragmatism

A lack of political will and pragmatism amplified the poor adoption and operationalisation of IWRM, a poorly performing economy and fleeing donor agencies resulted in less funding for water-related project. Political shenanigans created an imbalance that resulted in two forms of water i.e., water as an economic good vs. water as a social good [52]. <u>Manzungu</u> [53] argued post-colonial Zimbabwe continuously failed to develop a peoples-oriented water reform policy. In a bid to correct historical wrongs by availing subsidised water to the vulnerable and support the new social order, the initiative goes against the neo-liberalism approach that defines the “water as an economic good” [54] which is a founding principle of IWRM.

##### 4.3.1.2 South Africa

The water governance issue in the National Water Act of 1998 (NWA) aligns with the IWRM principles sought to provide equitable water distribution by re-writing the colonial disparities [55]. The drafting process was a multi-stakeholder and intersectoral activity that brought in international consultancies. Notable IWRM drivers were Department of International Development – UK (DFID), Danish Danida, and Deustsche Gesellschaft fur Zusammernarbeit (GIZ). The DFID was instrumental in water reform allocation law whilst the GIZ and Danida were active in experimental work in the catchments [56].

###### (i) Radical Innovation

Water redistribution in South Africa has been fraught with political and technical issues, for example, the Water Allocation Reform of 2003 failed to reconcile the apartheid disparity hence the equity component of IWRM was compromised. IWRM suffered another shock caused by the governing party when they introduced radical innovations that sought to shift from the socialist to neoliberal water resource use approach. The radical innovation through the government benefited the large-scale commercial farmers at the expense of the black smallholder farming community [55]. The radical innovation hindered policy diffusion from the Catchment Management Authority (CMA) and the large-scale commercial farmers to the smallholder irrigation schemes thus destabilising the IWRM adoption status.

###### (ii) Policy Misinterpretation

The shift from Integrated Catchment Management (ICM) to IWRM hindered the operationalisation and diffusion of the IWRM practice [57]. Despite acknowledging the “integration” researchers argued that the word lacked a clear-cut definition thus failing to establish a common ground for water’s multi-purpose use [55]. For maximised adoption of a practice, incremental innovation is required, which was Danida’s agenda in the quest to drive IWRM in South Africa. According to <u>Wehn and Montalvo</u> [58] incremental innovation “is characterised by marginal changes and occurs in mature circumstances, building and upgrading existing knowledge and skills”. IWRM diffusion encountered obstacles partly because the government could not ensure policy cascading.

###### (iii) Land Reform

Land reform in South Africa is characterised by (a) redistribution which seeks to transfer land from the white minority on a willing buyer willing seller basis, (b) restitution which rights the discriminatory 1913 land laws that saw natives evicted from their ancestral land, and (c) land tenure that provides tenure to the occupants of the homelands. This new pattern created a new breed of smallholder farmers that are, more often than not, excluded from diffusion and water governance channels [59]. In addition, researchers argue that a farm once owned by one white farmer is owned by multiple landowners with different cultural backgrounds and, more often than not, IWRM policy approach is met with resistance [60]. Another challenge posed by multicultural water users is the interpretation and translation of innovations.

###### (iv) Wet Water and Paper Water

To foster the water as an economic good aspect of IWRM the licensing system was enacted. The phenomenon was described by van Koppen (2012) as paper water precedes water, thus the disadvantaged black smallholder farmers could not afford paper water which consequently limits access to water. The licensing system can be interpreted as stifling the smallholder sector and hence negative attitudes develop and hinder policy adoption. Another issue that negatively impacted adoption was that issuing a license was subject to farmers possessing storage facilities. The smallholder farmers lack resources hence the requirement for obtaining a license excluded the small players in favour of the large-scale commercial farmers. This consequently maintains the historically skewed status-quo, where “big players” keep winning.

<u>Van Koppen</u> [61] and <u>Denby, Movik</u> [62] argue the shift from local water rights system to state-based water system have created bottlenecks making it hard for smallholder farmers to obtain “paper water” and subsequently “wet water”. The state-based system is characterised by bureaucracies and local norms are in perpetual change, hence denying the IWRM innovation policy approach stability efficiency.

#### 4.3.2 Prospect of IWRM in Southern Africa

The IWRM policy approach and practice in South Africa was government-driven whereas in Zimbabwe external donors were the main vehicles for diffusion. For both countries, the water and land reform agenda has a multiplicity of overlapping functionaries; however, they are managed by separate government departments. The silo system at national level prevents effective innovation diffusion and distorts policy cascading at the local level.

Water affairs are politicised and often, the water reform policy fails to balance the Dublin’s principles which form the backbone of the IWRM innovation policy approach. Failure by national governments to address unequal water access created by former segregationist policies is perpetuated by the lack of balance between creating a new social order and recognising the “water as an economic good” principle

## 5 Systematic Comparison of Findings on East, West and Southern Africa

Data extracts from the respective regional analysis were formulated into theoretical candidate themes. The thematic analysis extracted recurring themes common to all the three regions. (Table 4). The analysis performed a data extraction exercise and formulated codes (Figure 3). Themes were then generated from the coded data extracts to create a thematic map.

**Figure 3.**
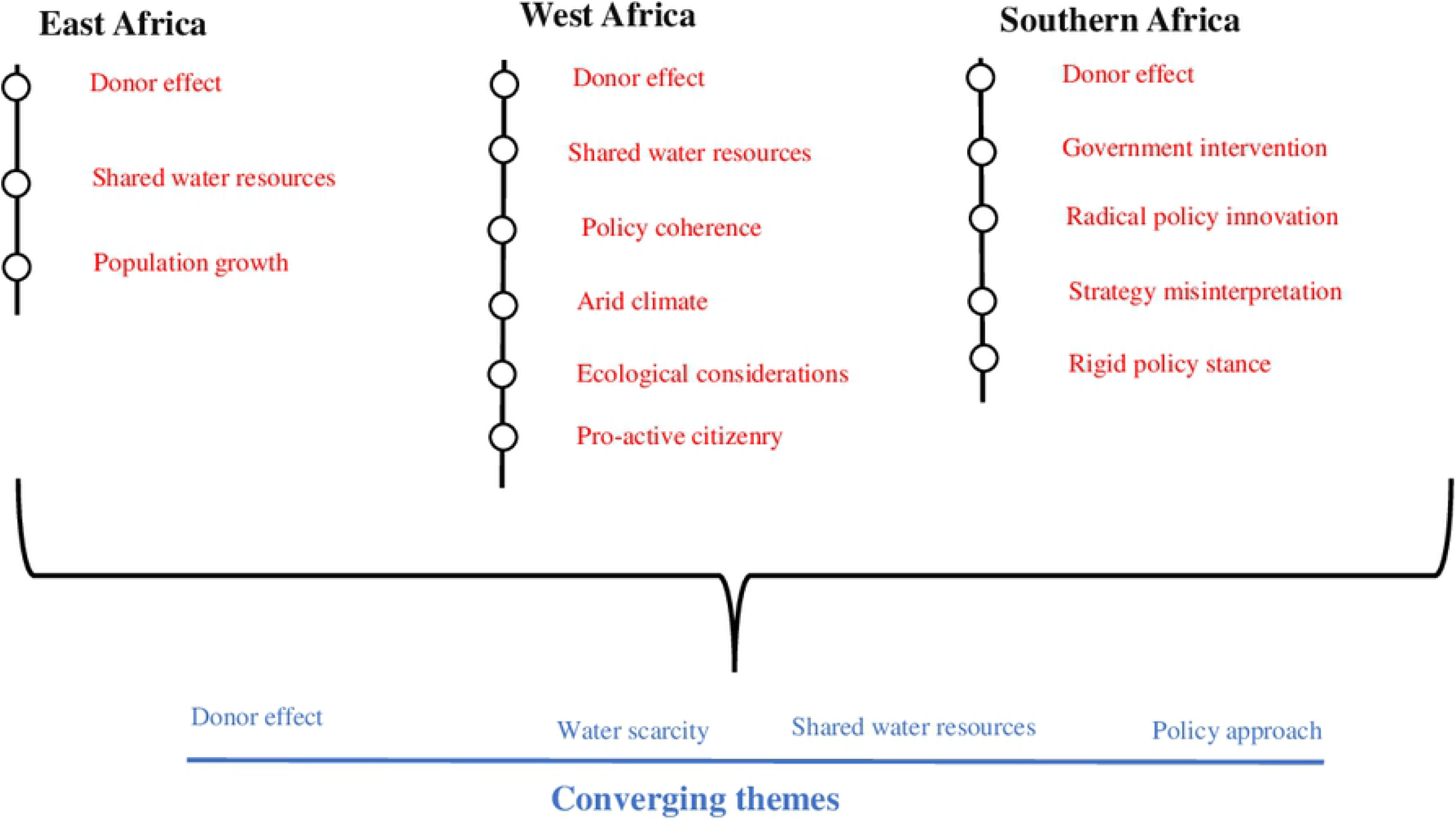
Extracted and converging themes from the coded data excerpts

**Table 4.**
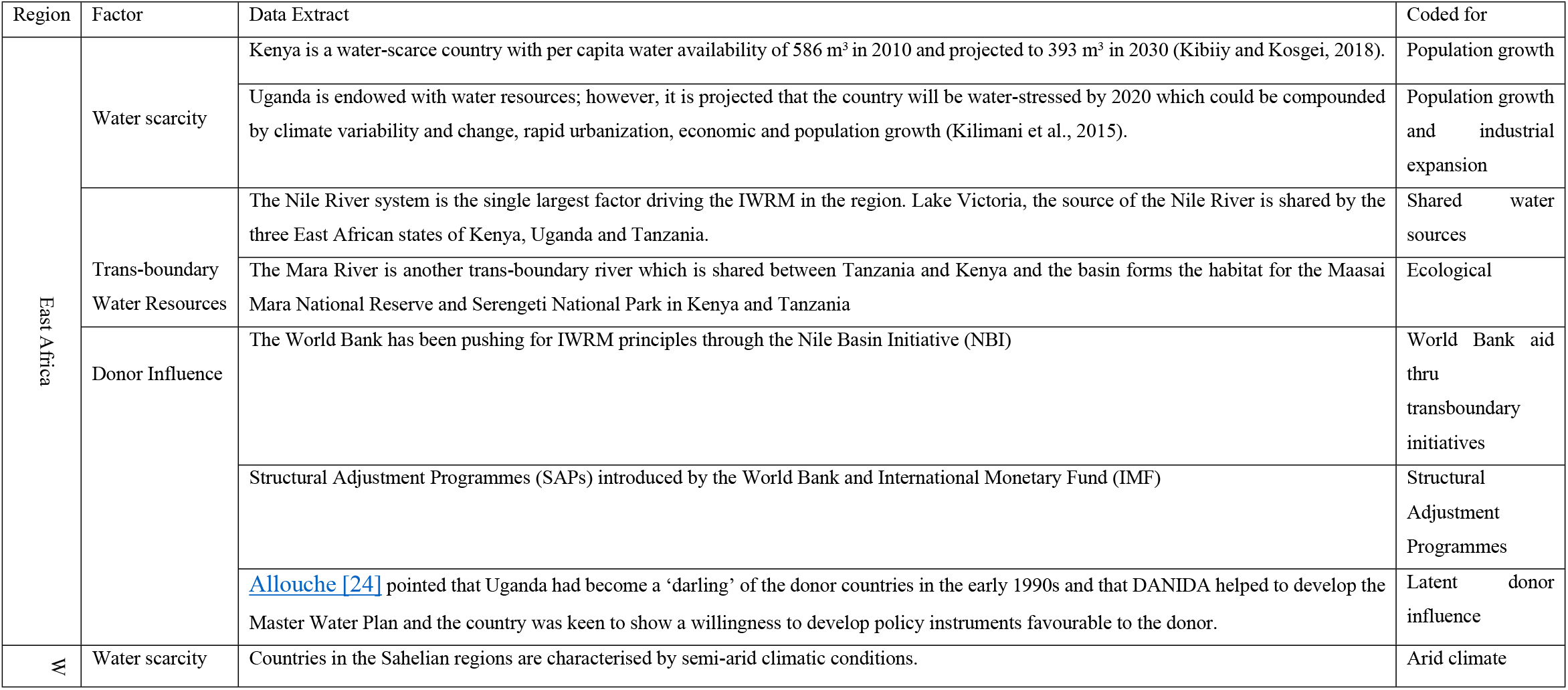
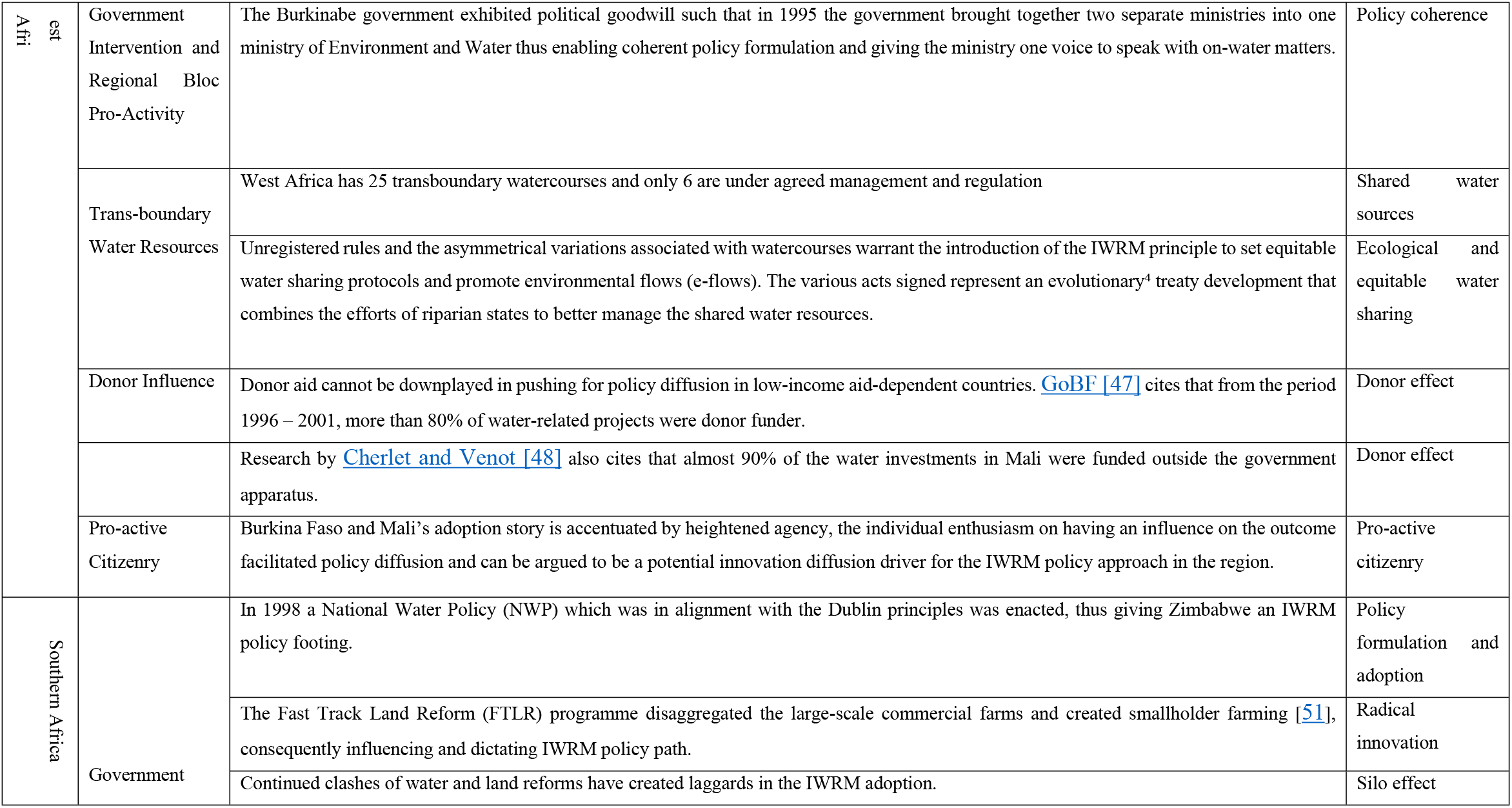
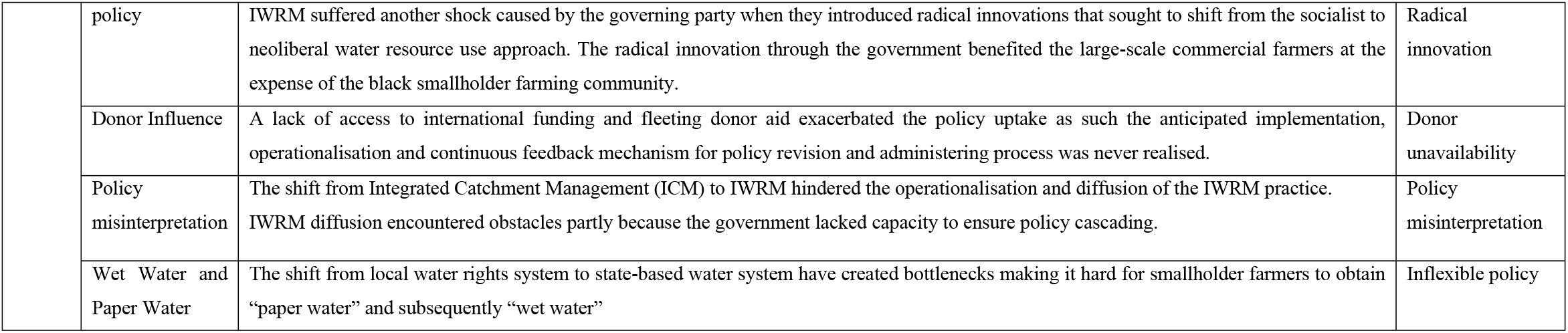
Data extracts with the applied codes.

### 5.1 Donor effect and Policy Approach

Donor activity invariably influenced the policy path that individual countries took. The three regions had significant support from donors to drive the IWRM strategy. Zimbabwe experienced a different fate. The political climate caused an exodus of donor support from the nation, which consequently caused a laggard. The absence of donor support was at the backdrop of the two formulated water acts namely National Water Act [63] and the Zimbabwe National Water Authority Act of 1998 [64], which were meant to promote equitable water provision amongst the population. This highlights the latent adoption of IWRM strategy. The 2008/2009 cholera outbreak raised alarm and facilitated the return of donor activity in Zimbabwe’s water sector. The availability of donor support motivated the redrafting of a water clause in the 2013 constitution that espoused the IWRM approach to water management [65].

Whilst <u>Mehta, Alba</u> [65] argue that South Africa enjoyed minimal donor support it cannot be downplayed how much donor influence impacted the IWRM strategy adoption. For instance, the Water Allocation Reform (WAR) was drafted with the aid of the UK Department of International Development. The WAR fundamentals are informed IWRM principles. The economic structural programmes spearheaded by The World Bank and the IMF were active in facilitating the diffusion of the IWRM strategy in Kenya and Uganda. Uganda made strides because of a long-standing relationship with donor nations. The Uganda – donor relationship dates back to early 1990 where Uganda was elected to be the NBI secretariat, this in itself evidence of commitment to water policy reform [5, 66].

### 5.2 Transboundary Water Resources

The Nile River system is the single largest factor driving the IWRM in the region. Lake Victoria, the source of the Nile River is shared by the three East African states of Kenya, Uganda and Tanzania. Irrigation schemes in Sudan and Egypt rely exclusively on the waters of River Nile and are therefore apprehensive of the actions of upstream states notably Ethiopia, Kenya, Uganda, Tanzania, Rwanda and Burundi. The source of contention is the asymmetrical water needs and allocation which was enshrined in the Sudan–Egypt treaty of 1959 [28]. All the riparian countries in the Nile basin have agricultural-based economies and thus irrigation is the cornerstone of food security [29]. Therefore, there was the need for the establishment of basin-wide co-operation which led to the formation of NBI in 1999 with a vision to achieve sustainable socio-economic development through the equitable utilisation of the Nile basin water resources [30]. In Eastern Africa, the Nile Basin Initiative (NBI) and the Lake Victoria Basin Commission (LVBC) plays a critical component in promoting the IWRM at regional level [17]. The LVBC is deeply intertwined with the East African Community (EAC) and thus has more political clout to implement policies regarding utilization of the Lake Victoria waters [67]. This, therefore, implies that for NBI to succeed, it must have a mandate and political goodwill from the member countries.

The conflicts around the utilization of the Nile water resources persists due to the treaty of 1959 which led to the signing of Cooperative Framework Agreement (CFA) by a number of the Nile basin countries, with the notable exceptions of Egypt, Sudan and South Sudan [68]. The CFA was signed between 2010 and 2011 and establishes the principle that each Nile Basin state has the right to use, within its territory, the waters of the Nile River Basin, and lays down some factors for determining equitable and reasonable utilization such as the contribution of each state to the Nile waters and the proportion of the drainage area [69]. The construction of the Grand Ethiopian Renaissance Dam has been a source of concern and conflict among the three riparian countries of Ethiopia, Sudan and Egypt Sudan [68]. The asymmetrical power relations (Egypt is the biggest economy) in the Nile Basin is a big hindrance to the co-operation among the riparian countries [70] and thus a threat to IWRM implementation in the shared watercourse. While Ethiopia is using its geographical power to negotiate for an equitable share in the Nile water resources, Egypt is utilizing both materials, bargaining and idealistic power to dominate the hydro politics in the region and thus the former can only succeed if it reinforces its geographical power with material power [71].

Therefore, IWRM implementation at the multi-national stage is complex but necessary to forestall regional conflicts and war. The necessity of co-operation rather than conflict in the Nile Basin is paramount due to the water availability constraints which is experienced by most countries in the region. The transboundary IWRM revolves around water-food-energy consensus where the needs of the riparian countries are sometimes contrasting, for example, Egypt and Sudan require the Nile waters for irrigation to feed their increasing population while Ethiopia requires the Nile waters for power generation to stimulate her economy. The upstream riparian States could use their bargaining power to foster co-operation and possibly force the hegemonic downstream riparian States into the equitable and sustainable use of Nile waters [72].

The SADC region has 13 major transboundary river basins (excluding the Nile and Congo) of Orange, Limpopo, Incomati, Okavango, Cunene, Cuvelai, Maputo, Buzi, Pungue, Save-Runde, Umbeluzi, Rovuma and Zambezi [73]. The Revised Protocol on Shared Watercourses is the key instrument for managing transboundary water resources in the SADC. The overall aim of the Protocol is to foster co-operation for judicious, sustainable and coordinated management, the protection and utilization of shared water resources [74].

<u>Ashton and Turton</u> [75] argue that the transboundary water issues in Southern Africa revolve around the key roles played by pivotal States and impacted States and their corresponding pivotal basins and impacted basins. In this case, pivotal States are riparian states with a high level of economic development (Botswana, Namibia, South Africa, and Zimbabwe) and a high degree of reliance on shared river basins for strategic sources of water supply while impacted States are riparian states (Angola, Lesotho, Malawi, Mozambique, Swaziland, Tanzania, and Zambia) that have a critical need for access to water from an international river basin that they share with a pivotal state, but appear to be unable to negotiate what they consider to be an equitable allocation of water and therefore, their future development dreams are impeded by the asymmetrical power dynamics with the pivotal states. Pivotal Basins (Orange, Incomati, and Limpopo) are international river basins that face closure but are also strategically important to anyone (or all) of the pivotal states by virtue of the range and magnitude of economic activity that they support. Impacted basins (Cunene, Maputo, Okavango, Cuvelai, Pungué, Save-Runde, and Zambezi) are those international river basins that are not yet approaching a point of closure, and which are strategically important for at least one of the riparian states with at least one pivotal State.

The transboundary co-operation under IWRM in Southern Africa is driven mainly by water scarcity which is predominant in most of the SADC countries which may imply the use of inter-basin transfers schemes [75]. Further, most of the water used for agriculture, industry and domestic are found within the international river basins [76] which calls for collaborative water management strategies. The tricky feature hindering the IWRM is the fact that States are reluctant to transfer power to River Basin Commissions [77]. Indeed most of the River Basin Organizations (RBO) in Southern region as the Zambezi Commission, the Okavango River Basin Commission, and the Orange-Sengu River Basin Commission have loose links with SADC and therefore lack the political clout to implement the policies governing the shared water resources [67]. Power asymmetry, like in Eastern Africa, is also a bottleneck in achieving equitable sharing of water resources as illustrated by the water transfer scheme involving Lesotho and South Africa [78]. The hydro-hegemonic South Africa is exercising control over any negotiations and agreements in the Orange-Senqu basin [79]. Limited data sharing among the riparian States is another challenge which affects water management in transboundary river basins e.g. in the Orange-Senqu basin [80].

West Africa has 25 transboundary watercourses and only 6 are under agreed management and regulation. The situation is compounded by the fact that 20 watercourses lack strategic riverbasin management instruments [45]. Unregistered rules and the asymmetrical variations associated with watercourses warrant the introduction of the IWRM principle to set equitable water sharing protocols and promote environmental flows (e-flows). The various acts signed represent an evolutionary^5^ treaty development that combines the efforts of riparian states to better manage the shared water resources. Water Resources Coordination Centre (WRCC) was established in 2004 to implement an integrated water resource management in West Africa and to ensure regional coordination of water resource related policies and activities [81].

The Niger River basin covers 9 Countries of Benin, Burkina, Cameroon, Chad, Côte d’Ivoire, Guinea, Mali, Niger and Nigeria. The Niger River Basin Authority (NBA) was established to promote co-operation among the member countries and to ensure basin-wide integrated development in all fields through the development of its resources, notably in the fields of energy, water resources, agriculture, livestock, forestry exploitation, transport and communication and industry [82]. The Shared Vision and Sustainable Development Action Programme (SDAP) was developed to enhance co-operation and sharing benefits from the resources of River Niger [83]. The Niger Basin Water Charter together with the SDAP are key instruments which set out a general approach to basin development, an approach negotiated and accepted not only by all member states but also by other actors who utilize the basin resources [84].

The main agreement governing the transboundary water resource in River Senegal Basin is the Senegal River Development Organization, OMVS (Organisation pour la mise en valeur du fleuve Sénégal) with its core principle being the equitably shared benefits of the resources of the basin [84]. The IWRM in the Senegal River Basin is hampered by weak institutional structures and lack of protocol on how shared waters among the States as well as conflicting national and regional interests [85, 86]. The Senegal River Basin, being situated in the Sudan-Sahelian region, is faced by the threat of climate change which affects water availability [86] The Senegal River Basin States have high risks of political instability.

In general, IWRM in West Africa is hampered by weak institutional structures.

## 6 Summary and Conclusion

Africa is a laboratory of IWRM produced varied aggregated outcomes. The outcomes were directly linked to various national socio-economic development agendas; thus, the IWRM policy took a multiplicity of paths. In East Africa, Kenya is still recovering from the devolved system of government to the County system which created new transboundary sectors with the country. Water scarcity, trans-boundary water resource and donor aid played a critical role in driving the IWRM policy approach in East and West Africa. Heightened agency and institutional integration allowed the diffusion of IWRM policy approach in rural West Africa.

Southern Africa’s adoption has been fraught with hindrances, for instance, land reform affected the effective adoption and operationalisation of the IWRM policy approach. Southern Africa has a unique political landscape that is closely tied to water reform policies thus governments are struggling to strike a balance between the new social order and the “water as an economic good” aspect of the IWRM policy. The silo effect demonstrated by the two southern African countries in question had negative effects on IWRM policy diffusion. Parallel land reform and water reform policy formulation and implementation emerged as a major hindrance for full adoption and implementation of IWRM. It is also worth noting that Zimbabwe’s case brought to light that water sector reforms cannot be pinned down to policy and political will, but a deep colonial history influenced the adoption and performance of the IWRM strategy.

For the future, IWRM policy approach can be implemented in Africa and the continent has the potential to implement and adopt the practice. Endowed with a significant number of water bodies, Africa must adopt the IWRM strategy for maximising regional cooperation and subsequent economic gains.

## Footnotes

1 Key: Substantiated (+++); partially substantiated (++); unsubstantiated (+) after Thomas, KM, et al., (2019) *SoE = Strength of Evidence

2 Evolutionary treaties can be classified as incremental innovation.

3 UNICEF administered a USD 83 million budget between 2013 & 2016. USD 53 million was meant for urban areas and the balance for rural development (Manzungu and Derman, 2016).

4 Evolutionary treaties can be classified as incremental innovation.

5 Evolutionary treaties can be classified as incremental innovation.

